# Molecular hydrogen is an overlooked energy source for marine bacteria

**DOI:** 10.1101/2022.01.29.478295

**Authors:** Rachael Lappan, Guy Shelley, Zahra F. Islam, Pok Man Leung, Scott Lockwood, Philipp A. Nauer, Thanavit Jirapanjawat, Ya-Jou Chen, Adam J. Kessler, Timothy J. Williams, Ricardo Cavicchioli, Federico Baltar, Perran L.M. Cook, Sergio E. Morales, Chris Greening

## Abstract

Molecular hydrogen (H_2_) and carbon monoxide (CO) are supersaturated in seawater relative to the atmosphere and hence are readily accessible energy sources for marine microbial communities. Yet while marine CO oxidation is well-described, it is unknown whether seawater communities consume H_2_. Here we integrated genome-resolved metagenomics, biogeochemistry, thermodynamic modelling, and culture-based analysis to profile H_2_ and CO oxidation by marine bacteria. Based on analysis of 14 surface water samples, collected from three locations spanning tropical to subantarctic fronts, three uptake hydrogenase classes are prevalent in seawater and encoded by major marine families such as Rhodobacteraceae, Flavobacteriaceae, and Sphingomonadaceae. However, they are less abundant and widespread than carbon monoxide dehydrogenases. Consistently, microbial communities in surface waters slowly consumed H_2_ and rapidly consumed CO at environmentally relevant concentrations, with H_2_ oxidation most active in subantarctic waters. The cell-specific power from these processes exceed bacterial maintenance requirements and, for H_2_, can likely sustain growth of bacteria with low energy requirements. Concordantly, we show that the polar ultramicrobacterium *Sphingopyxis alaskensis* grows mixotrophically on H_2_ by expressing a group 2a [NiFe]-hydrogenase, providing the first demonstration of atmospheric H_2_ oxidation by a marine bacterium. Based on TARA Oceans metagenomes, genes for trace gas oxidation are globally distributed and are fourfold more abundant in deep compared to surface waters, highlighting that trace gases are important energy sources especially in energy-limited waters. Altogether, these findings show H_2_ is a significant energy source for marine communities and suggest that trace gases influence the ecology and biogeochemistry of oceans globally.

## Introduction

Over the last decade, it has emerged that trace gases are major energy sources supporting the growth and survival of aerobic bacteria ^1^. Two trace gases, molecular hydrogen (H_2_) and carbon monoxide (CO), are particularly dependable substrates given their ubiquity, diffusibility, and energy yields ^2,3^. Bacteria oxidise these gases, including below atmospheric concentrations, using group 1 and 2 [NiFe]-hydrogenases and form I carbon monoxide dehydrogenases linked to aerobic respiratory chains ^4–9^. Trace gas oxidation enables diverse organoheterotrophic bacteria to survive long-term starvation for their preferred organic growth substrates ^10,11^. In addition, this process can support mixotrophic growth on various organic and inorganic energy sources ^10,12,13^. To date, bacteria from eight different phyla have been experimentally shown to consume H_2_ and CO at ambient levels ^7,12–19^, with numerous other bacteria encoding the determinants of this process ^9,20^. At the ecosystem scale, most bacteria in soil ecosystems harbour genes for trace gas oxidation and cell-specific rates of trace gas oxidation are theoretically sufficient to sustain their survival ^21,22^. However, given most of these studies have focused on soil environments or isolates, the wider significance of trace gas oxidation remains largely unexplored.

Trace gases are particularly relevant energy sources for oceanic bacteria given they are generally available at elevated concentrations related to the atmosphere, in contrast to most soils. Surface layers of the world’s oceans are generally supersaturated with H_2_ and CO, typically by 2- to 5-fold (up to 15-fold) and 20- to 200-fold (up to 2000-fold) relative to the atmosphere respectively ^23–26^. As a result, oceans contribute to net atmospheric emissions of these gases ^27,28^. CO is mainly produced through photochemical oxidation of dissolved organic matter ^29^, whereas H_2_ is primarily produced by cyanobacterial nitrogen fixation ^30^. High concentrations of H_2_ are also produced during fermentation in hypoxic sediments which can diffuse into the overlying water column, especially in coastal waters ^31^. For unresolved reasons, the distributions of these gases vary with latitude and exhibit opposite trends: while dissolved CO is highly supersaturated in polar waters, H_2_ is often undersaturated ^32–37^. These variations likely reflect differences in the relative rates of trace gas production and consumption in different climates.

Oceanic microbial communities have long been known to consume CO, though their capacity to use H_2_ has not been systematically evaluated ^38^. Approximately a quarter of bacterial cells in oceanic surface waters encode CO dehydrogenases in surface waters and these span a wide range of taxa, including the globally abundant family Rhodobacteraceae (marine *Roseobacter* clade) ^9,39–42^. Building on observations made for soil oxidation, CO oxidation potentially enhances the long-term survival of marine bacteria during periods of organic carbon starvation ^9^; consistently, culture-based studies indicate CO does not influence growth of marine isolates, but the enzymes responsible are strongly upregulated during starvation ^43–46^. While aerobic and anaerobic H_2_ oxidation has been extensively described by benthic and hydrothermal vent communities ^47–49^, to date no studies have shown whether pelagic bacterial communities can use this gas. Several surveys have detected potential H_2_-oxidising hydrogenases in seawater samples and isolates ^9,20,49,50^. While Cyanobacteria are well-reported to oxidise H_2_, including marine isolates such as *Trichodesmium*, this process is thought to be limited to the endogenous recycling of H_2_ produced by the nitrogenase reaction ^51,52^.

In this study, we addressed these knowledge gaps by investigating the mediators, rates, and potential roles of H_2_ and CO oxidation by marine bacteria. To do so, we performed side-by-side metagenomic and biogeochemical profiling of 14 samples collected from a temperate oceanic transect, a temperate coastal transect, and a tropical island, and also tested the capacity of three axenic marine bacterial isolates to aerobically consume atmospheric H_2_. We provide definitive ecosystem-scale and culture-based evidence that H_2_ is a relevant energy source for marine bacteria, though is only used by a small proportion of community members in contrast to CO.

## Results and Discussion

### Marine microbial communities slowly consume H_2_ and rapidly consume CO

We measured *in situ* concentrations and *ex situ* oxidation rates of H_2_ and CO in 14 surface seawater samples using an ultra-sensitive gas chromatograph. The samples were collected from three locations **(Fig. S1)**: an oceanic transect spanning neritic, subtropical, and subantarctic front waters (Munida Transect off New Zealand coast; n = 8; **Fig. S2**); a temperate urban bay (Port Phillip Bay, Australia; n = 4); and a tropical coral island (Heron Island, Australia; n = 2). In line with global trends, both gases were supersaturated relative to the atmosphere in all samples. H_2_ was supersaturated by 5.4-, 4.8- and 12.4-fold respectively in the oceanic transect (2.0 ± 1.2 nM), the temperate bay (1.8 ± 0.26 nM), and, as previously reported ^53^, the tropical island (4.6 ± 0.3 nM). CO was moderately supersaturated in the oceanic transect (5.2-fold; 0.36 nM ± 0.07 nM), but highly oversaturated in both the temperate bay (123-fold; 8.5 ± 1.7 nM) and tropical island (118-fold; 8.2 ± 0.93 nM).

Microbial oxidation of trace gases was detected in all but one of the collected samples during *ex situ* incubations **(Fig. 1)**. For the temperate bay, H_2_ and CO were consumed in water samples collected from the coast, intermediary zone, and bay centre **(Fig. 1a)**. Based on *in situ* gas concentrations, bulk oxidation rates of CO were 18-fold faster than H_2_ (*p* < 0.0001) **(Table S1)**. Bulk oxidation rates did not significantly differ between the surface microlayer (i.e. the 1 mm interface between the atmosphere and ocean) and underlying waters. H_2_ and CO oxidation was also evident in surface microlayer and underlying seawater samples collected from the tropical island **(Fig. S3)**. We similarly observed rapid CO and slower H_2_ consumption across the multi-front oceanic transect, though unexpectedly these activities were mutually exclusive. Net CO oxidation occurred throughout the coastal and subtropical waters, but was negligible in subantarctic waters. Conversely, net H_2_ oxidation only occurred in the subantarctic waters **(Fig. 1b)**. These divergent oxidation rates may help to explain the contrasting concentrations of H_2_ and CO in global seawater ^32–37^, though wider sampling and *in situ* assays would be required to confirm this. It should be noted that these measurements likely underestimate rates and overestimate thresholds of H_2_ oxidation given there will still be underlying endogenous production of H_2_ through nitrogen fixation during the incubations. Nevertheless, they provide the first definitive report of H_2_ oxidation in marine water columns.

**Figure 1.**
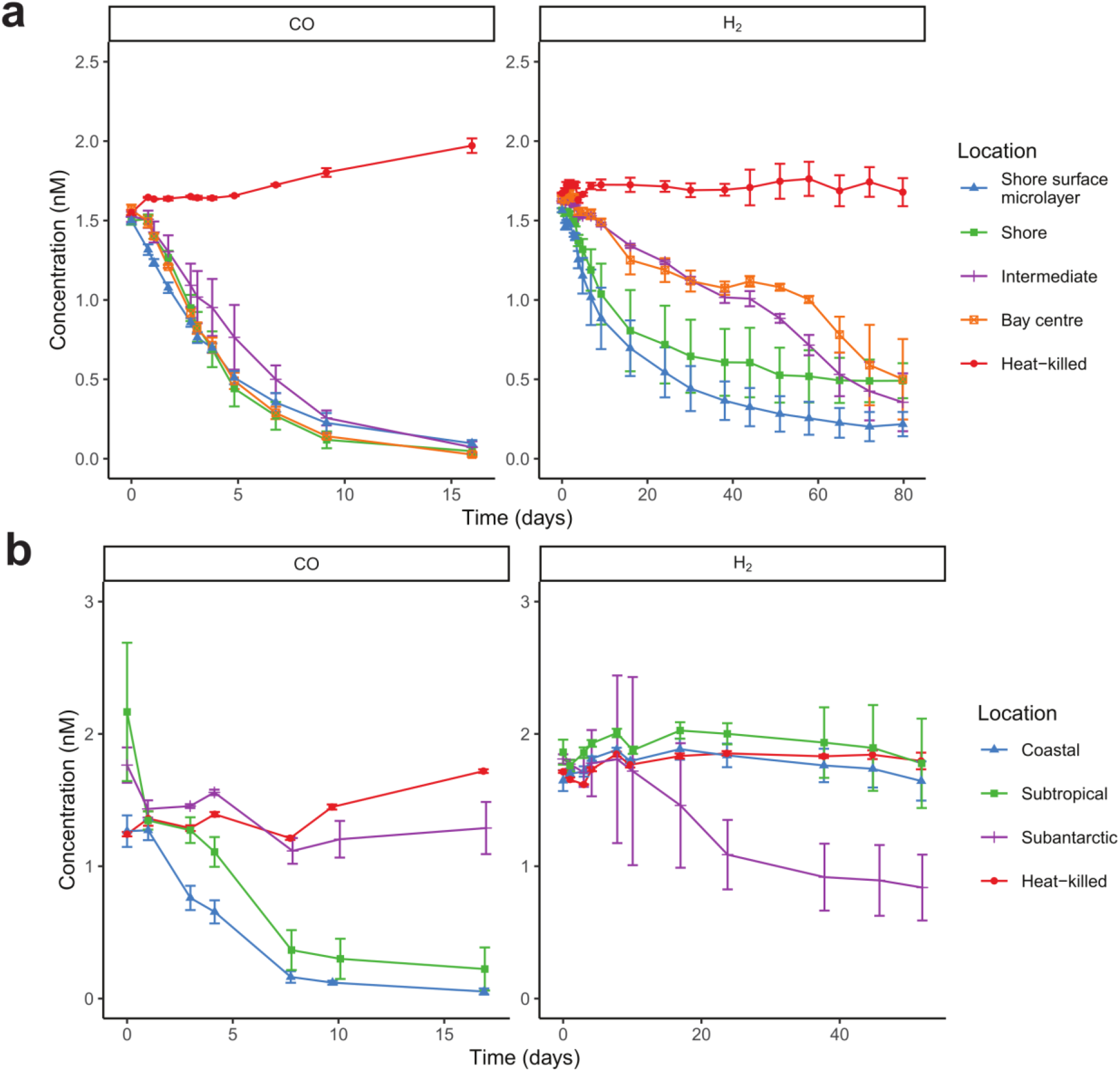
*Ex situ* oxidation of CO and H_2_ by seawater communities. Results are shown for **(a)** four samples in a transect of Port Phillip Bay, Victoria, Australia and **(b)** eight samples in the Munida transect off the coast of Otago, New Zealand. Each 120 mL sealed serum vial contained 60 mL of native seawater samples incubated in a 60 mL ambient air headspace supplemented with ~2.5 ppmv H_2_ or CO. At each timepoint, the mixing ratio of each gas in the headspace of each vial was measured on a gas chromatograph and converted to dissolved gas concentrations (nM).

### Marine microbial communities encode enzymes for both CO and H_2_ oxidation

To better understand the basis of these activities, we sequenced metagenomes of the 14 samples **(Table S2 & S3)**, which were assembled and binned into 110 medium- and high-quality metagenome-assembled genomes (MAGs) **(Table S4)**. We used homology-based searches to determine the abundance of 50 metabolic marker genes in the metagenomic reads **(Table S3)**, assemblies **(Table S5)**, and MAGs **(Table S4)**. In common with other surface seawater communities ^54^, analysis of community composition **(Fig. S4; Fig. 2b)** and metabolic genes **(Fig. 2a & 2b)** suggests most bacteria present are capable of organoheterotrophy, phototrophy, and aerobic respiration. The capacity for aerobic CO oxidation was moderately abundant. Approximately 12% of bacterial and archaeal cells encoded the *coxL* gene (encoding the catalytic subunit of the form I CO dehydrogenase), though relative abundance decreased from an average of 25% in the temperate bay where CO oxidation was highly active to 5.1% in subantarctic waters where CO oxidation was negligible **(Fig. 2a; Fig. 1)**. In contrast, H_2_ oxidation was a rare trait: the catalytic subunit of aerobic H_2_-uptake [NiFe]-hydrogenases was encoded by an average of 1.1% of bacteria across the samples **(Fig. 2a; Table S3)**. Hydrogenase abundance peaked in the tropical island samples (average 3.5%), but declined to 0.11% in the neritic and subtropical samples from the oceanic transect **(Fig. 2a)**, in line with the contrasting H_2_ oxidation rates between these samples **(Fig. 1; Fig. S3)**. Abundance of H_2_- and CO-oxidising bacteria strongly predicted oxidation rates of each gas (*R*^2^ of 0.55 and 0.88 respectively) **(Fig. S5)**, though it is likely that repression of gene expression contributes to the negligible activities of some samples.

**Figure 2.**
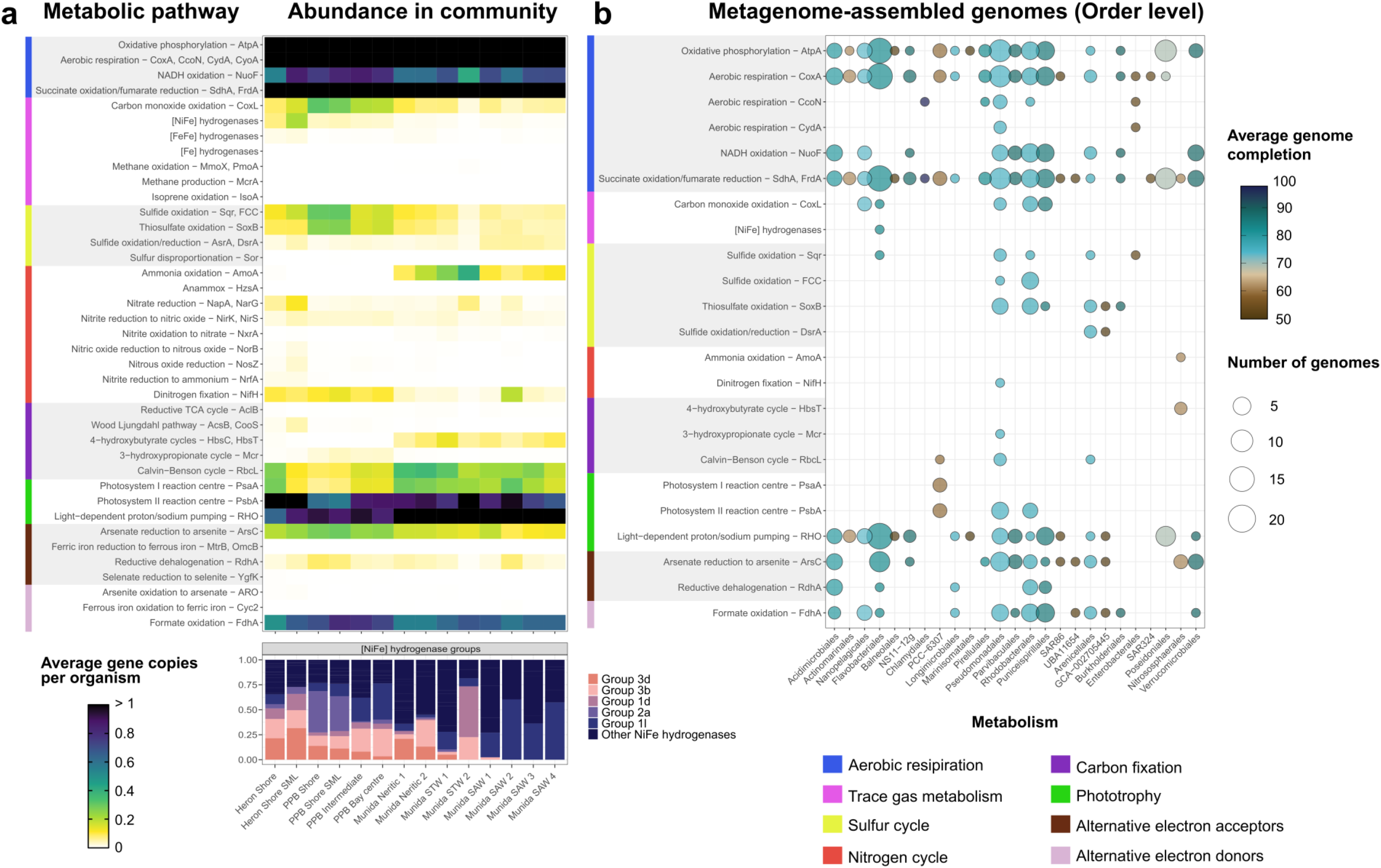
Abundance and distribution of metabolic genes encoded by marine communities. **(a)** Heatmap showing the abundance of metabolic marker genes in the metagenomic short reads across the three sampling locations and 14 samples. A homology-based search was used to calculate the relative abundance of marker genes as average gene copies per organism (abundance relative to a set of universal single-copy marker genes), and is equivalent to the estimated proportion of the community encoding a given gene as a single copy. Where multiple marker genes are listed, values are summed. The bottom panel shows the hydrogenase subgroups present in each sample. **(b)** Bubble plot showing metabolic potential of the 110 metagenome-assembled genomes (MAGs). MAGs are summarised at order level, with the size of the circle corresponding to the number of genomes in that order with a given gene, and the colour reflecting the percentage of genome completeness. Marker genes are omitted that were not detected in any MAG.

Based on the metagenome data, the ability to oxidise CO was consistently a more common and widespread metabolic strategy than the oxidation of H_2_. Diverse form I CO dehydrogenase genes, mostly affiliating with the recently defined proteobacterial, actinobacterial, and mixed 1 clades of the enzyme ^9^, were detected in the metagenomic short reads and assemblies **(Table S3 & S5)**. This diversity was reflected in the assembled metagenomes, with the gene encoded by 13 MAGs (12%) from the families Rhodobacteraceae, Flavobacterales UA16, Litoricolaceae, Puniceispirillaceae, and Nanopelagicales S36-B12 **(Fig. 2b; Table S4)**. All but two of these MAGs also encoded the genes for energy-converting rhodopsins or photosystem II, indicating they can harvest energy concurrently or alternately from both CO and light, in support of previous culture-based findings ^46^. While most of these MAGs are predicted to be obligate heterotrophs, two Rhodobacteraceae MAGs also encoded type IA ribulose 1,5-bisphosphate carboxylase/oxygenase (RuBisCO) and hence are theoretically capable of carboxydotrophic growth **(Fig. 2b; Table S4)**. These findings support previous inferences that habitat generalists in marine waters depend on metabolic flexibility and use dissolved CO to enhance growth or survival ^40,55^.

Based on the metagenomic reads and assemblies, three H_2_-uptake hydrogenases likely account for the observed oxidation activities **(Fig. 2a)**. The group 1l [NiFe]-hydrogenase, recently discovered in Antarctic saline soils ^7^, was encoded in most samples and was the sole uptake hydrogenase in the subantarctic samples **(Fig. 2a)**; reads and assemblies for this enzyme closely affiliated with the hydrogenases of the reference genomes from the Rhodobacteraceae isolates *Pseudaestuariivita atlantica* and *Marinovum algicola* **(Table S3 & S5)**. The group 2a [NiFe]-hydrogenase, which supports mixotrophic growth in diverse bacteria ^12^, was relatively abundant in the temperate bay and tropical island samples **(Fig. 2a)**. We recovered one MAG encoding this enzyme, from the genus UBA3478 within the Flavobacteriaceae **(Fig. 2b; Table S4)**, as well as unbinned contigs closely related to the hydrogenase of this MAG and *Sphingopyxis alaskensis* **(Table S5)**. The group 1d [NiFe]-hydrogenase, associated with aerobic hydrogenotrophic growth in diverse species ^20,47^, was also abundant in the surface microlayer samples **(Fig. 2a)**. Altogether, this assortment of hydrogenases suggests a small proportion of marine bacteria (1.1%) from at least three dominant marine families (Flavobacteriaceae, Rhodobacteraceae, Sphingomonadaceae) have established a stable niche by using an abundant substrate to support growth and potentially persistence. The sole hydrogenase-encoding MAG also encoded genes for succinate oxidation and rhodopsin-dependent light harvesting, suggesting H_2_ oxidation either supports mixotrophic growth or is a facultative trait.

### H_2_ can theoretically support survival and likely growth of marine bacteria

We used thermodynamic modelling to determine the amount of power (i.e. W per cell) generated based on the observed rates of trace gas oxidation **(Fig. 1; Table S1)** and predicted number of trace gas oxidisers **(Fig. 2a; Table S1)** in the sampled waters. This analysis was limited to the samples where oxidation was observed and reliable cell counts are available. On average, oxidation of the measured *in situ* concentrations of CO and H_2_ yields 7.2 × 10^−16^ W and 5.8 × 10^−14^ W per cell **(Fig. 3)**. The power derived from both trace gases is well within the range to sustain maintenance functions of bacteria, based on measurements of mostly copiotrophic isolates ^56,57^. Marine H_2_ oxidisers gain a particularly high amount of power by oxidising a relatively exclusive substrate at rapid cell-specific rates.

**Figure 3.**
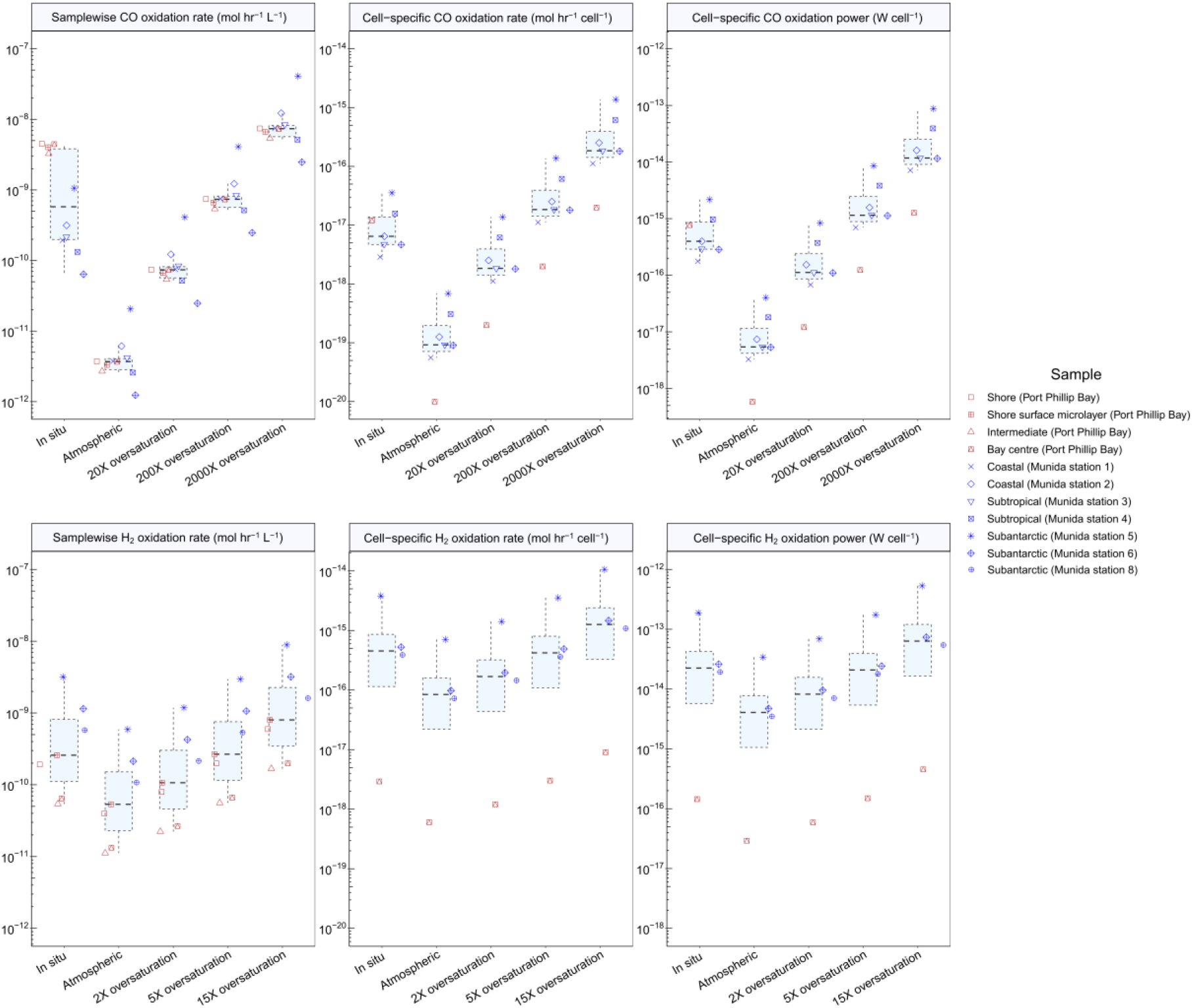
Thermodynamic modelling of H_2_ and CO oxidation by marine bacteria. The results show the bulk oxidation rates (left), oxidation rates per cell (middle), and power yields per cell (right) for **(a)** CO oxidation and **(b)** H_2_ oxidation. This analysis was only performed for samples where trace gas oxidation was measurable and cell-specific power was only calculated for samples where prokaryotic cell counts are available. Rates and power are shown based on CO and H_2_ concentrations at a range of environmentally relevant concentrations.

It is theoretically possible that the power derived from H_2_ oxidation supports growth. The cell-specific power generated for the sample with the most active H_2_ oxidisers (5.4 × 10^−13^ W; from the first subantarctic station) is below the growth requirements of most copiotrophic isolates, but likely sufficient to enable growth of the exceptionally small bacteria (ultramicrobacteria) that thrive in oligotrophic oceanic waters ^58^. Moreover, these power per cell calculations are likely underestimates given they do not account for any internal cycling of trace gases and assume all cells are equally active, and substantially increase when H_2_ and CO become transiently highly elevated over space and time as depicted in **Fig. 3**. Altogether, these considerations make it even more plausible that a small proportion of bacteria in oceans can grow using H_2_. By predominantly relying on energy derived from H_2_ oxidation, marine bacteria could potentially allocate most organic carbon for biosynthesis rather than respiration, i.e. adopting a predominantly lithoheterotrophic lifestyle.

### A marine isolate uses atmospheric H_2_ to supplement mixotrophic growth

To gain a better understanding of the mediators and roles of marine H_2_ oxidation, we investigated H_2_ uptake by three marine isolates encoding uptake hydrogenases. Two strains, *Robiginitalea biformata* DSM-15991 (Flavobacteriaceae) ^59^ and *Marinovum algicola* FF3 (Rhodobacteraceae) ^60^, did not substantially consume H_2_ over a three-week period across a range of conditions despite encoding group 1l [NiFe]-hydrogenases. It is unclear if hydrogenases have become non-functional in these fast-growing laboratory-adapted isolates or if they are instead only active under very specific conditions. *Sphingopyxis alaskensis* RB2256 (Spingomonadaceae) ^61,62^, which encodes a plasmid-borne group 2a [NiFe]-hydrogenase, aerobically consumed H_2_ in a first-order kinetic process to sub-atmospheric levels **(Fig. 4)**. Abundant in oligotrophic polar waters, *S. alaskensis* requires minimal resources to replicate given it forms extremely small cells (<0.1 μm^3^) and has a streamlined genome ^62–65^. Previously thought to be an obligate organoheterotroph ^66^, the discovery that this oligotrophic ultramicrobacterium ^67^ uses an abundant reduced gas as an energy source further rationalises its ecological success. This is the first report of atmospheric H_2_ oxidation by a marine bacterium.

**Figure 4.**
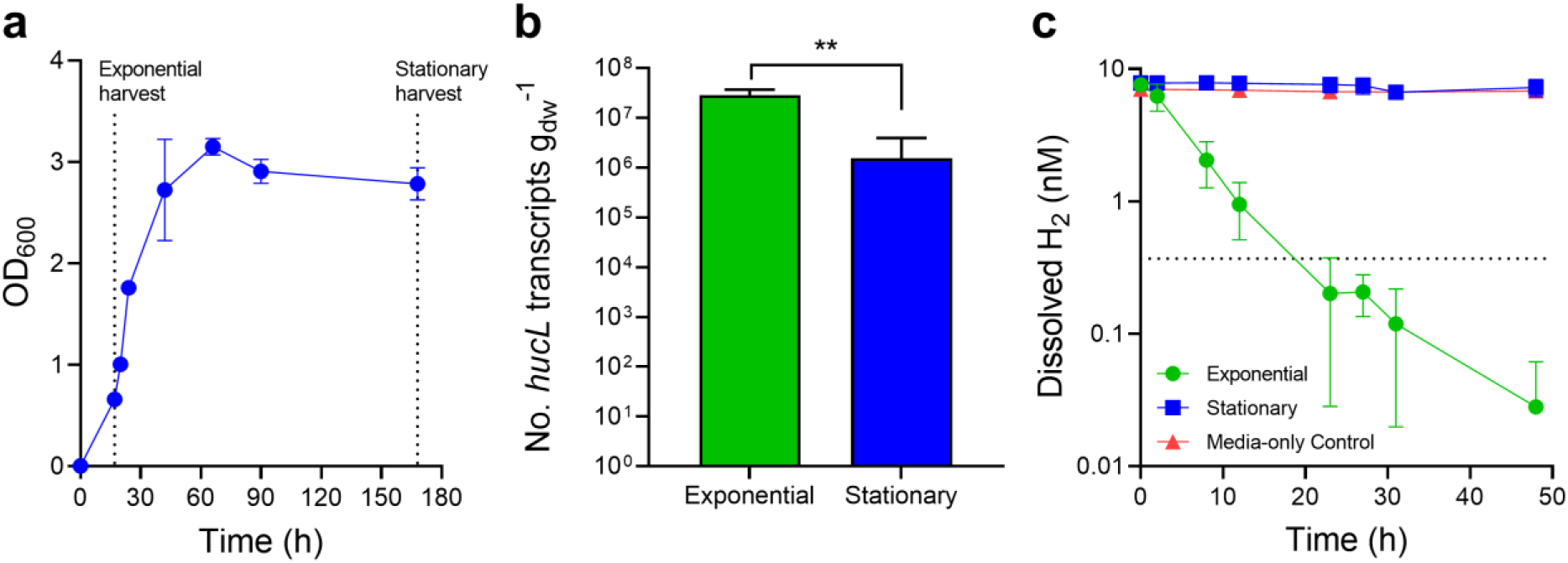
Hydrogenase expression and activity of *Sphingopyxis alaskensis*. **(a)** Growth curve of *S. alaskensis* grown on Difco 2216 Marine Broth. Cultures were tested for gas consumption and harvested for qPCR in exponential phase (17 h, OD600 = 0.66) and stationary phase (168 h, four days post-ODmax). **(b)** Number of transcripts of the group 2a [NiFe]-hydrogenase large subunit gene (*hucL*; locus Sala_3198), as measured by qRT-PCR, in exponential and stationary phase cultures of *S. alaskensis*. Error bars show standard deviations of three biological replicates (averaged from two technical duplicates) per condition. Values denoted by asterisks are statistically significant based on an unpaired t-test (*p* < 0.01). **(c)** H_2_ oxidation by exponential and stationary phase cultures of *Sphingopyxis alaskensis*. Error bars show the standard deviations of three biological replicates, with media-only vials monitored as negative controls. Dotted lines show the atmospheric concentration of hydrogen (0.53 ppmv).

We then determined whether *S. alaskensis* uses H_2_ oxidation primarily to support mixotrophic growth or survival. Expression levels of its hydrogenase large subunit gene (*hucL*) were quantified by qRT-PCR. Under ambient conditions, this gene was expressed at significantly higher levels (*p* = 0.006) during aerobic growth on organic carbon sources (mid-exponential phase; av. 2.9 × 10^7^ copies per gdw) than during survival (four days in stationary phase; av. 1.5 × 10^6^ copies per gdw; *p* = 0.006) **(Fig. 4a & 4b)**. This expression pattern is similar to other organisms possessing a group 2a [NiFe]-hydrogenase ^12^ and is antithetical to that of the groups 1h and 1l [NiFe]-hydrogenases that are typically induced by starvation ^7,14,16,68^. The activity of the hydrogenase was monitored under the same two conditions by monitoring depletion of headspace H_2_ mixing ratios over time by gas chromatography. H_2_ was rapidly oxidised by exponentially growing cultures to sub-atmospheric concentrations within a period of 30 hours, whereas negligible consumption was observed for stationary phase cultures **(Fig. 4c)**. Together, these findings suggest that *S. alaskensis* can grow mixotrophically in marine waters by simultaneously consuming dissolved H_2_ with available organic substrates. These findings align closely with that observed for other organisms harbouring group 2a [NiFe]-hydrogenases ^12,13^ and supports the inferences from thermodynamic modelling **(Fig. 3)** that H_2_ likely supports growth of some marine bacteria.

### Genes for trace gas oxidation are globally distributed and increase in relative abundance with ocean depth

We gained a global perspective on the distribution of the genes for H_2_ and CO oxidation by searching the metagenomes of the TARA Oceans dataset. Similarly to our metagenomes, H_2_-uptake hydrogenases (predominantly from groups 1d, 1l, and 2a) were quite rare in surface waters (encoded by 1.2% bacteria and archaea) and form I CO dehydrogenases were common (encoded by 15%). These genes were observed in samples spanning all four oceans, as well as the Red Sea and Mediterranean Sea **(Fig. 5)**. Importantly, we also observed that these genes increased in relative abundance by approximately fourfold in metagenomes from mesopelagic (200 to 1000 m deep) compared to surface waters (*p* < 0.0001). This pattern was consistent across sites in the Atlantic, Indian, Pacific, and Southern Oceans. These findings justify future studies comparing the rates and power associated with H_2_ and CO oxidation in deep compared to surface waters.

**Figure 5.**
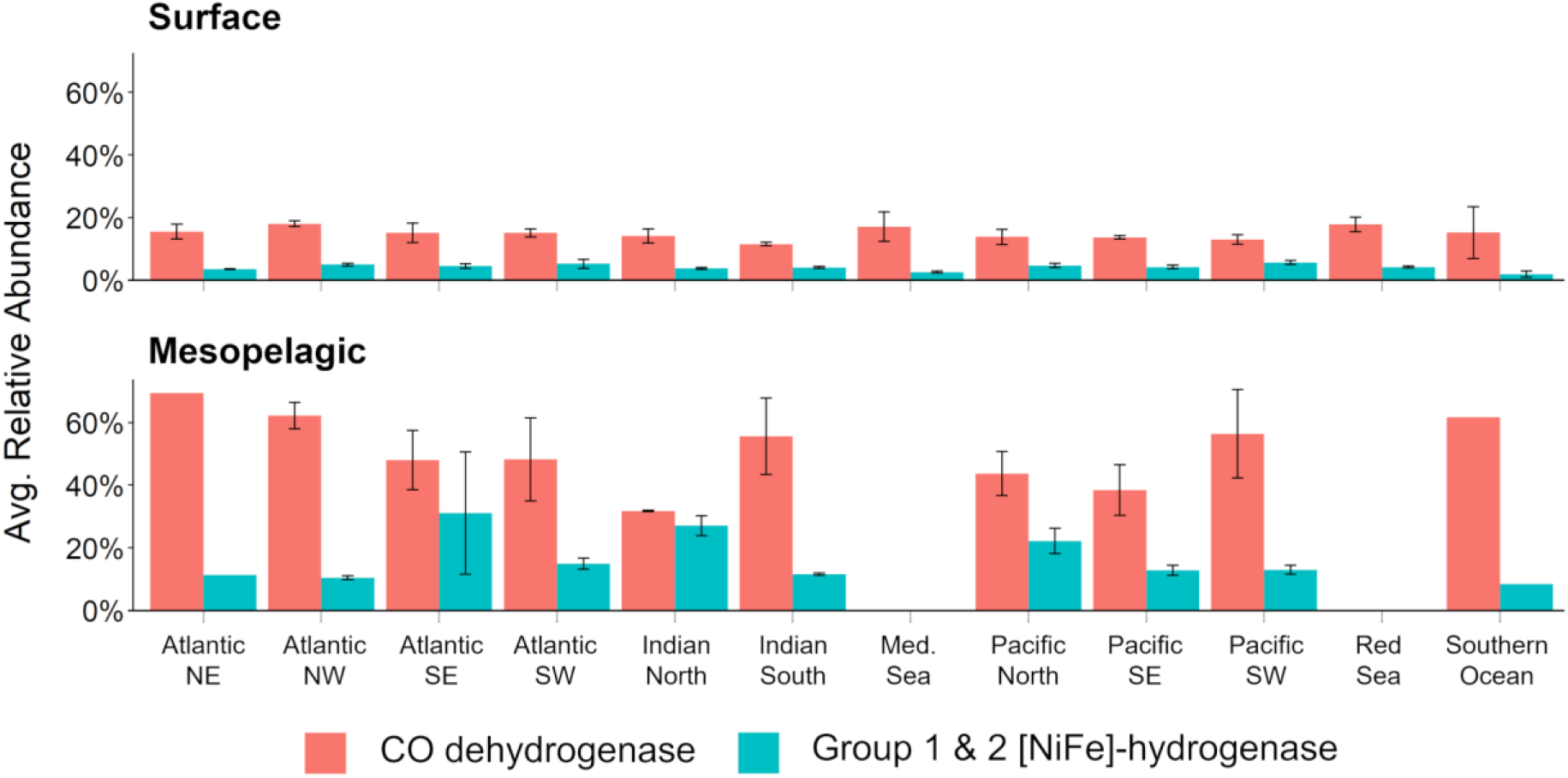
Relative abundance of CO and H_2_ oxidation genes in TARA Oceans metagenomes. The relative abundance of the catalytic subunit genes of the carbon monoxide dehydrogenase (CoxL) and group 1 & 2 [NiFe]-hydrogenases (inc. HucL and HylL) was normalised to a set of single-copy genes and averaged across all samples at a location. Locations are sorted by oceanic region (NE = Northeast, NW = Northwest, SE = Southeast, SW = Southwest, Med. = Mediterranean). No mesopelagic samples were sequenced for the Mediterranean Sea or Red Sea.

## Conclusions

Through an integrative approach, we provide the first demonstration that H_2_ is an important energy source for seawater communities. The biogeochemical, metagenomic, and thermodynamic modelling analyses together suggest that H_2_ is oxidised by a small proportion of community members, but at sufficiently fast cell-specific rates to enable mixotrophic growth. These findings are supported by experimental observations that the ultramicrobacterium *Sphingopyxis alaskensis* consumes H_2_ during heterotrophic growth. Marine bacteria likely gain a major competitive advantage from being able to consume this abundant, diffusible, high-energy gas. H_2_-oxidising marine microorganisms are globally distributed, though the activity-based measurements suggest complex controls on their activity and suggest they may be particularly active in low-chlorophyll waters. In contrast, our findings support that CO oxidation is a widespread trait that enhances the flexibility of habitat generalists ^39,40^, especially in high-chlorophyll waters. At the biogeochemical scale, our findings indicate marine bacteria mitigate atmospheric H_2_ emissions ^28^ and potentially account for undersaturation of H_2_ in Antarctic waters ^37^.

Yet a major enigma remains. H_2_ and CO are among the most dependable energy sources in the sea given their relatively high concentrations and energy yields. So why do relatively few bacteria harness them? By comparison, soils are net sinks for these trace gases given the numerous bacteria present rapidly consume them ^21^. We propose the straightforward explanation that the resource investment required to make the metalloenzymes to harness these trace gases may not always be justified by the energy gained. In the acutely iron-limited ocean, hydrogenases (containing 12-13 Fe atoms per protomer ^20^) and to a lesser extent CO dehydrogenases (containing four Fe atoms per protomer ^69^) are a major investment. This trade-off is likely to be most pronounced in the surface ocean, where solar energy that can be harvested using minimal resources through energy-converting rhodopsins. However, the iron investment required to consume H_2_ and CO is likely to be justified in energy-limited waters where primary production is low. This is consistent with the observed enrichment of hydrogenases and CO dehydrogenases in metagenomes from mesopelagic waters, as well as increased H_2_ oxidation observed in subantarctic waters. Thus, oceans continue to be a net source of H_2_ and CO despite the importance of these energy sources for diverse marine bacteria.

## Materials and Methods

### Sample collection and characteristics

To determine the ability of marine microbial communities to oxidise trace gases, a total of 14 marine surface water samples were collected from three different locations **(Fig. S1)**. Eight samples were collected from across the Munida Microbial Observatory Time-Series transect (Otago, New Zealand) ^70^ on 23/07/2019, in calm weather, on the *RV Polaris II*. This marine transect begins off the coast of Otago, New Zealand and extends through neritic, subtropical, and subantarctic waters ^70^. Eight equidistant stations were sampled travelling east, ranging from approximately 15 km to 70 km from Taiaroa Head. At each station, water was collected at 1 m depth using Niskin bottles and stored in two 1 L autoclaved bottles. One bottle was reserved for DNA filtration and extraction, whereas the other was used for microcosm incubation experiments. The vessel measured changes in salinity and temperature to determine the boundaries of each water mass (**Fig. S2**).

Four samples were also collected from the temperate Port Phillip Bay at Carrum Beach (Victoria, Australia) on 20/03/2019 and two were collected from the tropical Heron Island (Queensland, Australia) on 9/7/2019. At both sites, near-shore surface microlayer (SML) and surface water samples were collected in the subtidal zone (water depth ca. 1 m). At Port Phillip Bay, two samples were also collected at 7.5 km and 15 km east of the mouth of the Patterson River, labelled ‘Intermediate’ and ‘Centre’ respectively. In all cases, surface water samples of 3 L were collected with a sterile Schott bottle from approximately 20 cm depth and aliquoted for microcosm incubation and DNA extraction. SML samples were collected using a manual glass-plate sampler of 1800 cm^2^ surface area ^71^. A total of 520 – 580 mL was collected in 150 – 155 dips, resulting in an average sampling thickness of 20 μm. For the SML samples, 180 mL was reserved for microcosm incubations with the remaining volume used for DNA extraction. From all transects, each sample reserved for DNA extraction was vacuum-filtered using 0.22 μm polycarbonate filters, which were stored until extraction at −80°C.

### Measurement of dissolved H_2_ and CO

Dissolved gases were also sampled *in situ* at each transect to measure dissolved concentrations of CO and H_2_. Serum vials (160 mL) were filled with seawater using a gas-tight tube, allowing approximately 300 mL to overflow. The vial was then sealed with a treated lab-grade butyl rubber stopper, avoiding the introduction of gas to the vial. An ultra-pure N_2_ headspace (20 mL) was introduced to the vial by concurrently removing 20 mL of liquid, using two gas-tight syringes. The vials were then shaken vigorously for 2 minutes before equilibration for 5 minutes to allow dissolved gases to enter the headspace. 17 mL of the headspace was then collected into a syringe flushed with N_2_ by returning the removed liquid to the vial, and 2 mL was purged to flush the stopcock and needle before injecting the remaining 15 mL into a N_2_-flushed and evacuated silicone-closed Exetainer ^72^ for storage. Exetainers were sealed with a stainless-steel bolt and O-ring and stored until measurement. H_2_ and CO concentrations in the Exetainers were analysed by gas chromatography using a pulse discharge helium ionisation detector (model TGA-6792-W-4U-2, Valco Instruments Company Inc.), as previously described ^16^, calibrated against standard CO and H_2_ gas mixtures of known concentrations.

### *Ex situ* activity assays

To determine the ability of these marine microbial communities to oxidise CO and H_2_, the seawater samples were incubated with these gases under laboratory conditions and their concentration over time was measured with gas chromatography. For each sample, triplicate microcosms were setup in which seawater was transferred into foil-insulated serum vials (60 mL seawater in 120 mL vials for Munida transect and Port Phillip Bay; 80 mL seawater in 160 mL vials for Heron Island) and sealed with treated lab-grade butyl rubber stoppers ^72^. For each sampling location, one set of triplicates was also autoclaved and used as a control. The ambient air headspace of each vial was spiked with H_2_ and CO so that they reached initial headspace mixing ratios of either 2 ppmv (Munida transect and Port Phillip Bay) or 10 ppmv (Heron Island). Microcosms were continuously agitated at 20°C on a shaker table at 100 rpm. For Munida and Port Phillip Bay samples, 1 mL samples were extracted daily from the headspace and their content was measured by gas chromatography as described above. For Heron Island samples, at each timepoint, 6 mL gas was extracted and stored in 12 mL UHP-He-flushed conventional Exetainers (2018) or pre-evacuated 3 mL silicone-sealed Exetainers ^72^.

### Calculation of dissolved gas concentrations

The dissolved concentrations of gases in seawater at equilibrium state and at 1 atmospheric pressure were calculated according to the Sechenov relation for mixed electrolyte solutions as described by Weisenberger & Schumpe (1996) ^73^:

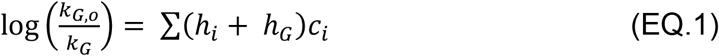

where *k*_*G,0*_ and *k*_*G*_ denote the gas solubility (or Henry’s law constant in equivalent) in water and the mixed electrolyte solution, respectively, *h*_*i*_ is a constant specific to the dissolved ion *i* (m^3^ kmol^−1^), *h*_*G*_ is a gas-specific parameter (m^3^ kmol^−1^), and *c*_*i*_ represents the concentration of the dissolved ion *i* in solution (kmol m^−3^). The gas-specific constant, *h*_*G*_, at temperature *T* (in K) follows the equation:

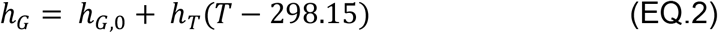

where *h*_*G,0*_ represents the value of *h*_*G*_ at 298.15 K and *h*_*T*_ is a gas-specific parameter for the temperature effect (m^3^ kmol^−1^ K^−1^). The gas solubility parameter *k*_*G,0*_ at temperature *T* follows combined Henry’s law and van’ t Hoff equation:

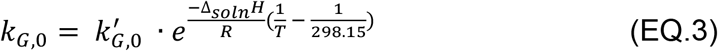

where 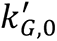 denotes Henry’s law constant of the gas at 298.15 K, Δ_*soln*_*H* is the enthalpy of solution and *R* is the ideal gas law constant.

The dissolved concentrations of gases at equilibrium with the headspace gas phase, at 1 atmospheric pressure and incubation temperature 20° C were calculated based on a mean seawater composition reported in Dickson and Goyet (1994) ^74^. The salinity correcting constants *h*_*i*_, *h*_*G,0*_, *h*_*T*_ were adopted from Weisenberger & Schumpe (1996) ^73^ while the temperature correcting constants 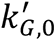 and 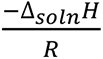 were obtained from Sander (2015) ^75^.

### Kinetic analysis and thermodynamic modelling

For kinetic analysis, measurement time points up to 30 days of incubation time were used. The gas consumption pattern was fitted with both an exponential model and a linear model. The former showed a lowest overall Akaike information criterion value for both H_2_ and CO consumption **(Table S1)**. As such, first order reaction rate constants were calculated and used for the kinetic modelling. In addition, only samples having at least two replicates with a positive rate constant were deemed to have a confident gas consumption. Bulk atmospheric gas oxidation rates for each sample were calculated with respect to the mean atmospheric mixing ratio of the corresponding trace gases (H_2_: 0.53 ppmv; CO: 0.09 ppmv; CH_4_: 1.9 ppmv). To estimate the cell-specific gas oxidation rate, the average direct cell count values reported for surface seawaters at Port Phillip Bay centre ^76^ and the eight stations along the Munida transect were used ^70,76^. Assuming all cells are viable and active, cell-specific gas oxidation rates were then inferred by dividing cell counts and the proportion of corresponding gas oxidisers from the metagenomic analysis.

To estimate the energetic contributions of H_2_ and CO oxidation to the corresponding marine trace gas oxidisers, we performed thermodynamics modelling to calculate their respective theoretical energy yields according to the first order kinetics of each sample estimated above. Power (Gibbs energy per unit time per cell), *P* follows the equation:

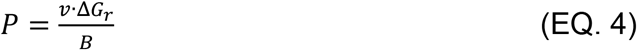

where *v* denotes the rate of substrate consumption per L of seawater (mol L^−1^ s^−1^) and *B* is the number of microbial cells (cells L^−1^) performing the reactions H_2_ + 0.5 O_2_ → H_2_O (dihydrogen oxidation) and CO + 0.5 O_2_ → CO_2_ (carbon monoxide oxidation). Δ*G*_*r*_ represents the Gibbs free energy of the reaction at the experimental conditions (J mol^−1^) and follows the equation:

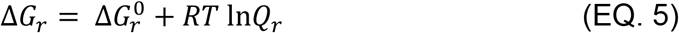

where 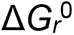 denotes the standard Gibbs free energy of the reaction, *Q*_*r*_ denotes the reaction quotient, *R* represents the ideal gas constant, and *T* represents temperature in Kelvin. Values of 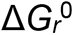 of the hydrogen oxidation and carbon monoxide oxidation were obtained from Thauer et al. (1977) ^77^. Values of *Q*_*r*_ for each reaction were calculated using:

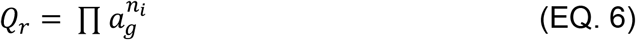

where *a*_*i*_ and *n*_i_ denote the dissolved concentration of the *i*^th^ species in seawater and the stoichiometric coefficient of the *i*^th^ species in the reaction of interest, respectively. Gibbs free energy for oxidation of hydrogen and carbon monoxide at atmospheric pressure and incubation temperature 20°C was calculated.

### Metagenomic sequencing and analysis

DNA was extracted from the sample filters using the DNeasy PowerSoil kit (QIAGEN) as per manufacturer’s instructions. Sample libraries, including an extraction blank control, were prepared with the Nextera XT DNA Sample Preparation Kit (Illumina) and sequenced on an Illumina NextSeq500 platform (2 × 151 bp) at the Australian Centre for Ecogenomics (University of Queensland). An average of 20,122,526 read pairs were generated per sample, with 827,868 read pairs sequenced in the negative control (**Table S2**). Raw metagenomic data was quality controlled with the BBTools suite v38.90 (https://sourceforge.net/projects/bbmap/), using BBDuk to remove the 151^st^ base, trim adapters, filter PhiX reads, trim the 3’ end at a quality threshold of 15 and discard reads below 50 bp in length. Reads detected in the extraction blank were additionally removed with BBMap v38.90, leaving a total of 97.7% of raw sample reads for further analysis. High quality short reads were then profiled for taxonomy by assembling and classifying 16S rRNA and 18S rRNA genes with PhyloFlash v3.4 ^78^. Short reads were assembled individually with metaSPAdes v3.14.1 ^79^ and collectively (all samples together, and by location) with MEGAHIT v1.2.9 ^80^. Coverage profiles for each contig were generated by mapping the short reads to the assemblies with BBMap v38.90 ^81^.

Genome binning was performed with MetaBAT2 v2.15.5 ^82^, MaxBin 2 v2.2.7 ^83^ and CONCOCT v1.1.0 ^84^ setting each tool to retain only contigs ≥ 2000 bp in length. For each assembly, resulting bins were dereplicated across binning tools with DAS_Tool v1.1.3 ^85^. All bins were refined with RefineM v0.1.2 ^86^ and consolidated into a final set of non-redundant metagenome-assembled-genomes (MAGs) at the default 99% average nucleotide identity using dRep v3.2.2 ^87^. The completeness, contamination and strain heterogeneity of each MAG was calculated with CheckM v1.1.3 ^88^, resulting in a total of 21 high-quality (> 90% completeness, < 5% contamination ^89^) and 89 medium-quality (> 50% completeness, < 10% contamination ^89^) MAGs. Taxonomy was assigned to each MAG with GTDB-Tk v1.6.0 ^90^ (using GTDB release 202) ^91^ and open reading frames were predicted from each MAG and additionally across all contigs (binned and unbinned) with Prodigal v2.6.3 ^92^. CoverM v0.6.1 (https://github.com/wwood/CoverM) “genome” was used to calculate the relative abundance of each MAG in each sample (--min-read-aligned-percent 0.75, --min-read-percent-identity 0.95, --min-covered-fraction 0) and the mean read coverage per MAG across the dataset (-m mean, --min-covered-fraction 0).

High quality short reads and predicted proteins from assemblies MAGs underwent metabolic annotation using DIAMOND v2.0.9 (--max-target-seqs 1, --max-hsps 1) ^93^ for alignment against a custom set of 50 metabolic marker protein databases. The marker proteins (https://doi.org/10.26180/c.5230745) cover the major pathways for aerobic and anaerobic respiration, energy conservation from organic and inorganic compounds, carbon fixation, nitrogen fixation, and phototrophy ^7^. Gene hits were filtered as follows: alignments were filtered to retain only those at least 40 amino acids in length (short read alignments) or with at least 80% query or 80% subject coverage (predicted MAG proteins). Alignments were further filtered by a minimum percentage identity score by gene: for short reads, this was 80% (PsaA), 75% (HbsT), 70% (PsbA, IsoA, AtpA, YgfK and ARO), 60% (CoxL, MmoA, AmoA, NxrA, RbcL, NuoF, FeFe hydrogenases and NiFe Group 4 hydrogenases), or 50% (all other genes). For predicted proteins, the same thresholds were used except for AtpA (60%), PsbA (60%), RdhA (45%), Cyc2 (35%) and RHO (30%). For short reads, gene abundance in the community was estimated as *average gene copies per organism* by dividing the abundance of the gene (in reads per kilobase million, RPKM) by the mean abundance of 14 universal single-copy ribosomal marker genes (in RPKM, obtained from the SingleM v0.13.2 package, https://github.com/wwood/singlem). For single-copy metabolic genes, this corresponds to the proportion of community members that encode the gene. A linear correlation analysis, performed in GraphPad Prism 9, was used to determine how metagenomic gene abundance correlated with *ex situ* H_2_ and CO oxidation rates.

### Culture-based growth and gas consumption analysis

Axenic cultures of three bacterial strains were analysed in this study: *Sphingopyxis alaskensis* (RB2256) ^61,62^ obtained from UNSW Sydney, *Robiginitalea biformata* DSM-15991 ^59^ imported from DSMZ, and *Marinovum algicola* FF3 (Rhodobacteraceae) ^60^ imported from DSMZ. Cultures were maintained in 120 mL glass serum vials containing a headspace of ambient air (H_2_ mixing ratio ~0.5 ppmv) sealed with treated lab-grade butyl rubber stoppers ^72^. Broth cultures of all three species were grown in 30 mL of Difco 2216 Marine Broth media and incubated at 30°C at an agitation speed of 150 rpm in a Ratek Orbital Mixer Incubator with access to natural day/night cycles. Growth was monitored by determining the optical density (OD_600_) of periodically sampled 1 mL extracts using an Eppendorf BioSpectrophotometer. The ability of the three cultures to oxidise H_2_ was measured with gas chromatography. Cultures in biological triplicate were opened, equilibrated with ambient air (1 h), and resealed. These re-aerated vials were then amended with H_2_ (via 1% v/v H_2_ in N_2_ gas cylinder, 99.999% pure) to achieve final headspace concentrations of ~10 ppmv. Headspace mixing ratios were measured immediately after closure and at regular intervals thereafter until the limit of quantification of the gas chromatograph was reached (42 ppbv H_2_). This analysis was performed for both exponential (OD_600_ 0.67 for *S. alaskensis*) and stationary phase cultures (~72 h post OD_max_ for *S. alaskensis*).

### Quantitative RT-PCR analysis

Quantitative reverse transcription PCR (qRT-PCR) was used to determine the expression levels of the group 2a [NiFe]-hydrogenase large subunit gene (*hucL*; locus Sala_3198) in *S. alaskensis* during growth and survival. For RNA extraction, triplicate 30 mL cultures of *S. alaskensis* were grown synchronously in 120 mL sealed serum vials. Cultures were grown to either exponential phase (OD600 0.67) or stationary phase (48 h post OD_max_ ~3.2). Cells were then quenched using a glycerol-saline solution (−20°C, 3:2 v/v), harvested by centrifugation (20,000 x *g*, 30 min, −9°C), resuspended in 1 mL cold 1:1 glycerol:saline solution (−20°C), and further centrifuged (20,000 x *g*, 30 min, −9°C). Briefly, resultant cell pellets were resuspended in 1 mL TRIzol Reagent (Thermo Fisher Scientific), mixed with 0.1 mm zircon beads (0.3 g), and subject to beat-beating (three 30 s on / 30 s off cycles, 5000 rpm) in a Precellys 24 homogenizer (Bertin Technologies) prior to centrifugation (12,000 x *g*, 10 min, 4°C). Total RNA was extracted using the phenol-chloroform method as per manufacturer’s instructions (TRIzol Reagent User Guide, Thermo Fisher Scientific) and resuspended in diethylpyrocarbonate-treated water. RNA was treated using the TURBO DNA-free kit (Thermo Fisher Scientific) as per manufacturer’s instructions. RNA concentration and purity were confirmed using a NanoDrop ND-1000 spectrophotometer.

cDNA was synthesised using a SuperScript III First-Strand Synthesis System kit for qRT-PCR (Thermo Fisher Scientific) with random hexamer primers, as per manufacturer’s instructions. Quantitative RT-PCR was performed using a LightCycler 480 SYBR Green I Master Mix (Roche) as per manufacturer’s instructions in 96-well plates and conducted in a QuantStudio 7 Flex Real-Time PCR System (Applied Biosystems). Primers were designed using Primer3 ^94^ to target the *hucL* gene (HucL_fw: AGCTACACAAACCCTCGACA; HucL_rvs: AGTCGATCATGAACAGGCCA) and the 16S rRNA gene as a housekeeping gene (16S_fwd: AACCCTCATCCCTAGTTGCC; 16S_rvs: GGTTAGAGCATTGCCTTCGG). Copy numbers for each gene were interpolated from standard curves of each gene created from threshold cycle (C_T_) values of amplicons that were serially diluted from 10^2^ to 10 copies (R^2^ > 0.98). Hydrogenase expression data was then normalised to the housekeeping in exponential phase. All biological triplicate samples, standards, and negative controls were run in technical duplicate. A student’s t-test in GraphPad Prism 9 was used to compare *hucL* expression levels between exponential and stationary phase.

## Footnotes

### Data availability statement

All raw metagenomes and metagenome-assembled genomes are deposited to the NCBI Sequence Read Archive under the BioProject accession number PRJNA801081. Metagenomics analysis scripts are publicly available at https://github.com/greeninglab/MarineOxidationManuscript

## Acknowledgements

This study was supported by ARC Discovery Project grants (DP180101762 and P210101595; both awarded to P.L.M.C. and C.G.), an ARC DECRA Fellowship (DE170100310; salary for C.G.), an NHMRC EL2 Fellowship (APP1178715; salary for C.G.). an Australian Government Research Training Stipend Scholarship (awarded to P.M.L.), Monash International Tuition Scholarships (awarded to P.M.L. and Y.J.C.), and Monash Postgraduate Publications Awards (awarded to Z.F.I. and Y.J.C.).

## Author contributions

C.G. conceived and supervised this study. C.G., G.S., S.E.M., P.L.M.C., R.L., and Z.I. designed experiments. G.S., S.L., S.E.M., P.A.N., Y.J.C., A.J.K., and P.L.M.C. contributed to field work. R.L., G.S., S.L., and C.G. contributed to metagenome analysis. G.S., P.M.L., P.A.N., and C.G. contributed to biogeochemical analysis. P.M.L., C.G., F.B., and P.M.L.C. contributed to thermodynamic modelling. Z.F.I., T.J., G.S., T.J.W., R.C., and C.G. contributed to culture-based work. C.G., R.L., and Z.F.I. wrote the paper with input from all authors. The authors declare no conflict of interest.

## References

1. Greening, C. & Grinter, R. Microbial oxidation of atmospheric trace gases. Nat. Rev. Microbiol. In press (2022).

2. Piché-Choquette, S. & Constant, P. Molecular hydrogen, a neglected key driver of soil biogeochemical processes. Appl. Environ. Microbiol. 85, e02418–18 (2019).

3. Greening, C., Islam, Z. F. & Bay, S. K. Hydrogen is a major lifeline for aerobic bacteria. Trends Microbiol. doi:10.1016/j.tim.2021.08.004 (2021).

4. Berney, M. & Cook, G. M. Unique flexibility in energy metabolism allows mycobacteria to combat starvation and hypoxia. PLoS One 5, e8614 (2010).

5. Greening, C., Berney, M., Hards, K., Cook, G. M. & Conrad, R. A soil actinobacterium scavenges atmospheric H_2_ using two membrane-associated, oxygen-dependent [NiFe] hydrogenases. Proc. Natl. Acad. Sci. U. S. A. 111, 4257–4261 (2014).

6. Myers, M. R. & King, G. M. Isolation and characterization of *Acidobacterium ailaaui* sp. nov., a novel member of Acidobacteria subdivision 1, from a geothermally heated Hawaiian microbial mat. Int. J. Syst. Evol. Microbiol. 66, 5328–5335 (2016).

7. Ortiz, M. et al. Multiple energy sources and metabolic strategies sustain microbial diversity in Antarctic desert soils. Proc. Natl. Acad. Sci. USA In revision (2021).

8. King, G. M. Molecular and culture-based analyses of aerobic carbon monoxide oxidizer diversity. Appl. Environ. Microbiol. 69, 7257–7265 (2003).

9. Cordero, P. R. F. et al. Atmospheric carbon monoxide oxidation is a widespread mechanism supporting microbial survival. ISME J. 13, 2868–2881 (2019).

10. Greening, C., Villas-Bôas, S. G., Robson, J. R., Berney, M. & Cook, G. M. The growth and survival of *Mycobacterium smegmatis* is enhanced by co-metabolism of atmospheric H_2_. PLoS One 9, e103034 (2014).

11. Liot, Q. & Constant, P. Breathing air to save energy – new insights into the ecophysiological role of high-affinity [NiFe]-hydrogenase in *Streptomyces avermitilis*. Microbiologyopen 5, 47–59 (2016).

12. Islam, Z. F. et al. A widely distributed hydrogenase oxidises atmospheric H_2_ during bacterial growth. ISME J. 14, 2649–2658 (2020).

13. Leung, P. M. et al. A nitrite-oxidising bacterium constitutively oxidises atmospheric H_2_. bioRxiv (2021).

14. Constant, P., Chowdhury, S. P., Pratscher, J. & Conrad, R. Streptomycetes contributing to atmospheric molecular hydrogen soil uptake are widespread and encode a putative high-affinity [NiFe]-hydrogenase. Environ. Microbiol. 12, 821–829 (2010).

15. Greening, C. et al. Persistence of the dominant soil phylum Acidobacteria by trace gas scavenging. Proc. Natl. Acad. Sci. U. S. A. 112, 10497–10502 (2015).

16. Islam, Z. F. et al. Two Chloroflexi classes independently evolved the ability to persist on atmospheric hydrogen and carbon monoxide. ISME J. 13, 1801–1813 (2019).

17. Schmitz, R. A. et al. The thermoacidophilic methanotroph *Methylacidiphilum fumariolicum* SolV oxidizes subatmospheric H_2_ with a high-affinity, membrane-associated [NiFe] hydrogenase. ISME J. 14, 1223–1232 (2020).

18. Hardy, K. R. & King, G. M. Enrichment of high-affinity CO oxidizers in Maine forest soil. Appl. Environ. Microbiol. 67, 3671–3676 (2001).

19. King, C. E. & King, G. M. Description of *Thermogemmatispora carboxidivorans* sp. nov., a carbon-monoxide-oxidizing member of the class Ktedonobacteria isolated from a geothermally heated biofilm, and analysis of carbon monoxide oxidation by members of the class Ktedonobacter. Int. J. Syst. Evol. Microbiol. 64, 1244–1251 (2014).

20. Greening, C. et al. Genomic and metagenomic surveys of hydrogenase distribution indicate H_2_ is a widely utilised energy source for microbial growth and survival. ISME J. 10, 761–777 (2016).

21. Bay, S. K. et al. Trace gas oxidizers are widespread and active members of soil microbial communities. Nat. Microbiol. 6, 246–256 (2021).

22. Xu, Y. et al. Genome-resolved metagenomics reveals how soil bacterial communities respond to elevated H_2_ availability. Soil Biol. Biochem. 163, 108464 (2021).

23. Schmidt, U. Molecular hydrogen in the atmosphere. Tellus 26, 78–90 (1974).

24. Walter, S. et al. Isotopic evidence for biogenic molecular hydrogen production in the Atlantic Ocean. Biogeosciences 13, 323–340 (2016).

25. Moore, R. M. et al. Extensive hydrogen supersaturations in the western South Atlantic Ocean suggest substantial underestimation of nitrogen fixation. J. Geophys. Res. Ocean. 119, 4340–4350 (2014).

26. Conte, L., Szopa, S., Séférian, R. & Bopp, L. The oceanic cycle of carbon monoxide and its emissions to the atmosphere. Biogeosciences 16, 881–902 (2019).

27. Khalil, M. A. K. & Rasmussen, R. A. The global cycle of carbon monoxide: Trends and mass balance. Chemosphere 20, 227–242 (1990).

28. Ehhalt, D. H. & Rohrer, F. The tropospheric cycle of H_2_: a critical review. Tellus, Ser. B Chem. Phys. Meteorol. 61, 500–535 (2009).

29. Miller, W. L. & Zepp, R. G. Photochemical production of dissolved inorganic carbon from terrestrial organic matter: Significance to the oceanic organic carbon cycle. Geophys. Res. Lett. 22, 417–420 (1995).

30. Moore, R. M., Punshon, S., Mahaffey, C. & Karl, D. The relationship between dissolved hydrogen and nitrogen fixation in ocean waters. Deep Sea Res. Part I Oceanogr. Res. Pap. 56, 1449–1458 (2009).

31. Kessler, A. J. et al. Bacterial fermentation and respiration processes are uncoupled in permeable sediments. Nat. Microbiol. 4, 1014–1023 (2019).

32. Swinnerton, J. W., Linnenbom, V. J. & Lamontagne, R. A. The ocean: a natural source of carbon monoxide. Science 167, 984–986 (1970).

33. Swinnerton, J. W. & Lamontagne, R. A. Carbon monoxide in the South Pacific Ocean. Tellus 26, 136–142 (1974).

34. Herr, F. L., Scranton, M. I. & Barger, W. R. Dissolved hydrogen in the Norwegian Sea: Mesoscale surface variability and deep-water distribution. Deep Sea Res. Part A. Oceanogr. Res. Pap. 28, 1001–1016 (1981).

35. Herr, F. L. Dissolved hydrogen in Eurasian Arctic waters. Tellus B 36, 55–66 (1984).

36. Conrad, R., Seiler, W., Bunse, G. & Giehl, H. Carbon monoxide in seawater (Atlantic Ocean). J. Geophys. Res. Ocean. 87, 8839–8852 (1982).

37. Conrad, R. & Seiler, W. Methane and hydrogen in seawater (Atlantic Ocean). Deep Sea Res. Part A. Oceanogr. Res. Pap. 35, 1903–1917 (1988).

38. Conrad, R. & Seiler, W. Photooxidative production and microbial consumption of carbon monoxide in seawater. FEMS Microbiol. Lett. 9, 61–64 (1980).

39. Tolli, J. D., Sievert, S. M. & Taylor, C. D. Unexpected diversity of bacteria capable of carbon monoxide oxidation in a coastal marine environment, and contribution of the *Roseobacter*-associated clade to total CO oxidation. Appl. Environ. Microbiol. 72, 1966–1973 (2006).

40. Mou, X., Sun, S., Edwards, R. A., Hodson, R. E. & Moran, M. A. Bacterial carbon processing by generalist species in the coastal ocean. Nature 451, 708–711 (2008).

41. Cunliffe, M. Correlating carbon monoxide oxidation with cox genes in the abundant marine *Roseobacter* clade. ISME J. 5, 685 (2011).

42. Royo-Llonch, M. et al. Compendium of 530 metagenome-assembled bacterial and archaeal genomes from the polar Arctic Ocean. Nat. Microbiol. (2021) doi:10.1038/s41564-021-00979-9.

43. Cunliffe, M. Physiological and metabolic effects of carbon monoxide oxidation in the model marine bacterioplankton *Ruegeria pomeroyi* DSS-3. Appl. Environ. Microbiol. 79, 738–740 (2013).

44. Christie-Oleza, J. A., Fernandez, B., Nogales, B., Bosch, R. & Armengaud, J. Proteomic insights into the lifestyle of an environmentally relevant marine bacterium. ISME J. 6, 124 (2012).

45. Muthusamy, S. et al. Comparative proteomics reveals signature metabolisms of exponentially growing and stationary phase marine bacteria. Environ. Microbiol. 19, 2301–2319 (2017).

46. Giebel, H.-A., Wolterink, M., Brinkhoff, T. & Simon, M. Complementary energy acquisition via aerobic anoxygenic photosynthesis and carbon monoxide oxidation by *Planktomarina temperata* of the *Roseobacter* group. FEMS Microbiol. Ecol. 95, fiz050 (2019).

47. Schwartz, E., Fritsch, J. & Friedrich, B. H_2_-metabolizing prokaryotes. (Springer Berlin Heidelberg, 2013).

48. Adam, N. & Perner, M. Microbially mediated hydrogen cycling in deep-sea hydrothermal vents. Front. Microbiol. 9, 2873 (2018).

49. Anantharaman, K., Breier, J. A., Sheik, C. S. & Dick, G. J. Evidence for hydrogen oxidation and metabolic plasticity in widespread deep-sea sulfur-oxidizing bacteria. Proc. Natl. Acad. Sci. 110, 330–335 (2013).

50. Barz, M. et al. Distribution analysis of hydrogenases in surface waters of marine and freshwater environments. PLoS One 5, e13846 (2010).

51. Eichner, M. J., Basu, S., Gledhill, M., de Beer, D. & Shaked, Y. Hydrogen dynamics in *Trichodesmium* colonies and their potential role in mineral iron acquisition. Front. Microbiol. 10, 1565 (2019).

52. Bothe, H., Schmitz, O., Yates, M. G. & Newton, W. E. Nitrogen fixation and hydrogen metabolism in cyanobacteria. Microbiol. Mol. Biol. Rev. 74, 529–551 (2010).

53. Nauer, P. A. et al. Pulses of labile carbon cause transient decoupling of fermentation and respiration in permeable sediments. bioRxiv In preparation (2022).

54. Sunagawa, S. et al. Structure and function of the global ocean microbiome. Science 348, 1261359 (2015).

55. Chen, Y. J. et al. Metabolic flexibility allows bacterial habitat generalists to become dominant in a frequently disturbed ecosystem. ISME J. 10.1038/s41396-021-00988-w (2021) doi:10.1101/2020.02.12.945220.

56. DeLong, J. P., Okie, J. G., Moses, M. E., Sibly, R. M. & Brown, J. H. Shifts in metabolic scaling, production, and efficiency across major evolutionary transitions of life. Proc. Natl. Acad. Sci. 107, 12941–12945 (2010).

57. LaRowe, D. E. & Amend, J. P. Power limits for microbial life. Front. Microbiol. 6, 718 (2015).

58. Lever, M. A. et al. Life under extreme energy limitation: A synthesis of laboratory- and field-based investigations. FEMS Microbiology Reviews vol. 39 688–728 (2015).

59. Cho, J.-C. & Giovannoni, S. J. *Robiginitalea biformata* gen. nov., sp. nov., a novel marine bacterium in the family Flavobacteriaceae with a higher G+ C content. Int. J. Syst. Evol. Microbiol. 54, 1101–1106 (2004).

60. Lafay, B. et al. *Roseobacter algicola* sp. nov., a new marine bacterium isolated from the phycosphere of the toxin-producing dinoflagellate Prorocentrum lima. Int. J. Syst. Evol. Microbiol. 45, 290–296 (1995).

61. Schut, F. et al. Isolation of typical marine bacteria by dilution culture: growth, maintenance, and characteristics of isolates under laboratory conditions. Appl. Environ. Microbiol. 59, 2150–2160 (1993).

62. Schut, F., Gottschal, J. C. & Prins, R. A. Isolation and characterisation of the marine ultramicrobacterium *Sphingomonas* sp. strain RB2256. FEMS Microbiol. Rev. 20, 363–369 (1997).

63. Vancanneyt, M. et al. *Sphingomonas alaskensis* sp. nov., a dominant bacterium from a marine oligotrophic environment. Int. J. Syst. Evol. Microbiol. 51, 73–79 (2001).

64. Eguchi, M. et al. *Sphingomonas alaskensis* strain AFO1, an abundant oligotrophic ultramicrobacterium from the North Pacific. Appl. Environ. Microbiol. 67, 4945–4954 (2001).

65. Cavicchioli, R., Ostrowski, M., Fegatella, F., Goodchild, A. & Guixa-Boixereu, N. Life under nutrient limitation in oligotrophic marine environments: an eco/physiological perspective of *Sphingopyxis alaskensis* (formerly *Sphingomonas alaskensis*). Microb. Ecol. 45, 203–217 (2003).

66. Williams, T. J., Ertan, H., Ting, L. & Cavicchioli, R. Carbon and nitrogen substrate utilization in the marine bacterium *Sphingopyxis alaskensis* strain RB2256. ISME J. 3, 1036–1052 (2009).

67. Lauro, F. M. et al. The genomic basis of trophic strategy in marine bacteria. Proc. Natl. Acad. Sci. 106, 15527–15533 (2009).

68. Berney, M., Greening, C., Hards, K., Collins, D. & Cook, G. M. Three different [NiFe] hydrogenases confer metabolic flexibility in the obligate aerobe Mycobacterium smegmatis. Environ. Microbiol. 16, 318–330 (2014).

69. Dobbek, H., Gremer, L., Meyer, O. & Huber, R. Crystal structure and mechanism of CO dehydrogenase, a molybdo iron-sulfur flavoprotein containing S-selanylcysteine. Proc. Natl. Acad. Sci. U. S. A. 96, 8884–8889 (1999).

70. Baltar, F., Stuck, E., Morales, S. & Currie, K. Bacterioplankton carbon cycling along the subtropical frontal zone off New Zealand. Prog. Oceanogr. 135, 168–175 (2015).

71. Cunliffe, M. & Wurl, O. Guide to best practices to study the ocean’s surface. Marine Biological Association of the United Kingdom for SCOR Plymouth, UK (Marine Biological Association of the United Kingdom for SCOR, 2014).

72. Nauer, P. A., Chiri, E., Jirapanjawat, T., Greening, C. & Cook, P. L. M. Technical note: Inexpensive modification of Exetainers for the reliable storage of trace-level hydrogen and carbon monoxide gas samples. Biogeosciences 18, (2021).

73. Weisenberger, S. & Schumpe, dan A. Estimation of gas solubilities in salt solutions at temperatures from 273 K to 363 K. AIChE J. 42, 298–300 (1996).

74. Dickson, A. G. & Goyet, C. Handbook of methods for the analysis of the various parameters of the carbon dioxide system in sea water; (Oak Ridge National Laboratory, 1994).

75. Sander, R. Compilation of Henry’s law constants (version 4.0) for water as solvent. Atmos. Chem. Phys. 15, 4399–4981 (2015).

76. Wenley, J. et al. Seasonal prokaryotic community linkages between surface and deep ocean water. Front. Mar. Sci. 8, 777 (2021).

77. Thauer, R. K., Jungermann, K. & Decker, K. Energy conservation in chemotrophic anaerobic bacteria. Bacteriol. Rev. 41, 100–180 (1977).

78. Gruber-Vodicka, H. R., Seah, B. K. B. & Pruesse, E. PhyloFlash: rapid small-subunit rRNA profiling and targeted assembly from metagenomes. mSystems 5, e00920–20 (2019).

79. Nurk, S., Meleshko, D., Korobeynikov, A. & Pevzner, P. A. metaSPAdes: a new versatile metagenomic assembler. Genome Res. 27, 824–834 (2017).

80. Li, D. H. et al. MEGAHIT v1.0: A fast and scalable metagenome assembler driven by advanced methodologies and community practices. Methods 102, 3–11 (2016).

81. Bushnell, B. BBMap: A Fast, Accurate, Splice-Aware Aligner. (2015).

82. Kang, D. et al. MetaBAT 2: an adaptive binning algorithm for robust and efficient genome reconstruction from metagenome assemblies. PeerJ 7, e7359 (2019).

83. Wu, Y.-W., Simmons, B. A. & Singer, S. W. MaxBin 2.0: an automated binning algorithm to recover genomes from multiple metagenomic datasets. Bioinformatics 32, 605–607 (2015).

84. Alneberg, J. et al. Binning metagenomic contigs by coverage and composition. Nat. Methods 11, 1144 (2014).

85. Sieber, C. M. K. et al. Recovery of genomes from metagenomes via a dereplication, aggregation and scoring strategy. Nat. Microbiol. 1 (2018).

86. Parks, D. H. et al. Recovery of nearly 8,000 metagenome-assembled genomes substantially expands the tree of life. Nat. Microbiol. 2, 1533 (2017).

87. Olm, M. R., Brown, C. T., Brooks, B. & Banfield, J. F. dRep: a tool for fast and accurate genomic comparisons that enables improved genome recovery from metagenomes through de-replication. ISME J. 11, 2864 (2017).

88. Parks, D. H., Imelfort, M., Skennerton, C. T., Hugenholtz, P. & Tyson, G. W. CheckM: assessing the quality of microbial genomes recovered from isolates, single cells, and metagenomes. Genome Res. 25, 1043–1055 (2015).

89. Bowers, R. M. et al. Minimum information about a single amplified genome (MISAG) and a metagenome-assembled genome (MIMAG) of bacteria and archaea. Nat. Biotechnol. 35, 725–731 (2017).

90. Chaumeil, P.-A., Mussig, A. J., Hugenholtz, P. & Parks, D. H. GTDB-Tk: a toolkit to classify genomes with the Genome Taxonomy Database. Bioinformatics 36, 1925–1927 (2020).

91. Parks, D. H. et al. A standardized bacterial taxonomy based on genome phylogeny substantially revises the tree of life. Nat. Biotechnol. 36, 996–1004 (2018).

92. Hyatt, D. et al. Prodigal: prokaryotic gene recognition and translation initiation site identification. BMC Bioinformatics 11, 119 (2010).

93. Buchfink, B., Xie, C. & Huson, D. H. Fast and sensitive protein alignment using DIAMOND. Nat. Methods 12, 59 (2014).

94. Untergasser, A. et al. Primer3—new capabilities and interfaces. Nucleic Acids Res. 40, e115–e115 (2012).

