## Supplementary information for "Molecular hydrogen is an overlooked energy source for marine bacteria"

**Table S1 (xlsx).** Trace gas oxidation rates and power calculations for Port Phillip Bay and Munida Transect samples.

**Table S2 (xlsx).** Sequencing statistics of the 15 sequenced metagenomes.

**Table S3 (xlsx).** Abundance of metabolic marker genes in the short read metagenomic data. This includes (a) the calculated average copies per organism for each marker gene, (b) average copies per organism for each hydrogenase subgroup, and (c) a list of short-read hits and their corresponding match in the database.

**Table S4 (xlsx).** Summary of metagenome-assembled-genomes (MAGs). This includes (a) descriptive information for each MAG, (b) relative abundance of each MAG per sample calculated by CoverM, (c) a summary of metabolic marker genes identified in each MAG, and (d) a full list of metabolic marker genes identified in the MAGs with alignment information and protein sequences.

**Table S5 (xlsx).** Marker genes identified in the assembled contigs. This lists the metabolic marker genes annotated across all contigs over 2000 bp (including both unbinned contigs and those that were assigned to a bin).

**Figure S1. Map of the sites sampled in this study.**

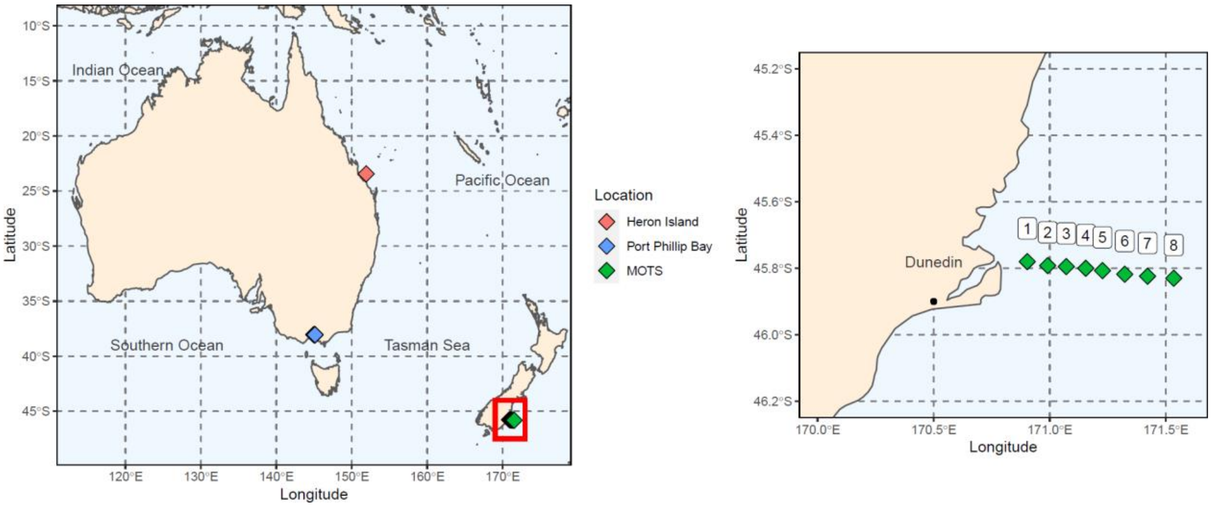

**Figure S2. Eight stations sampled in the Munida Observation Time Series.** The transect extends 65 km in an Eastern direction from Taieri Head, Otago. It spans neritic waters (NW; black; two stations), a transitional subtropical frontal zone (STW; yellow; two stations), and subantarctic waters (SAW; blue; four stations). Temperature and salinity data reflect the transitions between the three zones.

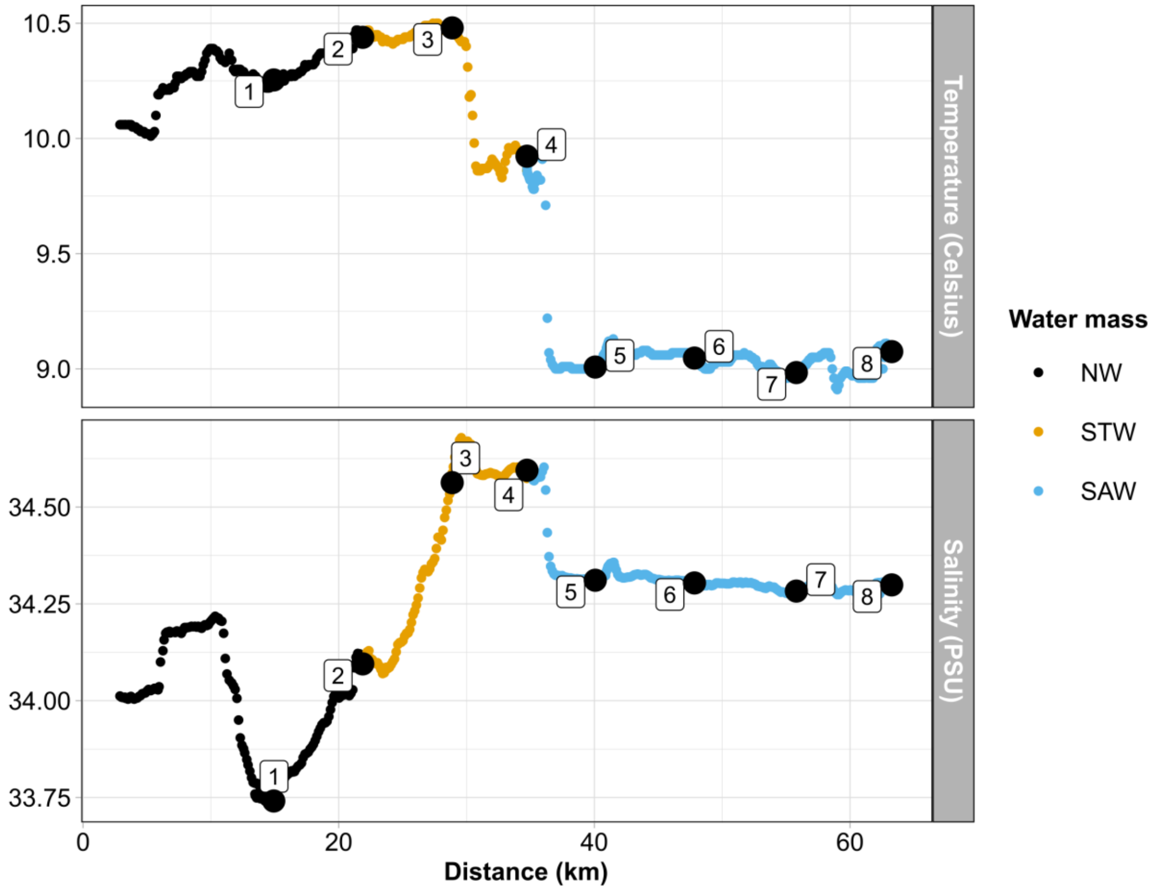

**Figure S3. Trace gas oxidation by marine microbial communities in Heron Island.** Samples were spiked with 10 ppm CO and $H_2$ and incubated in the dark. Relative concentrations were calculated by dividing concentrations at later times with the measured concentration at the start of each incubation.

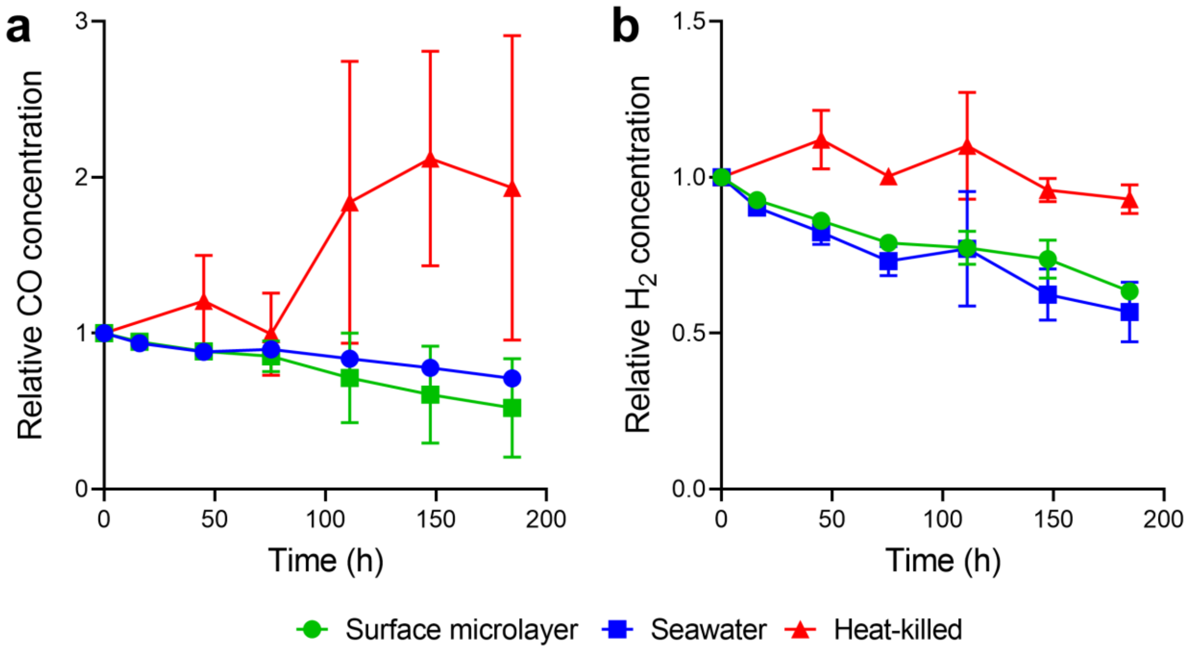

**Figure S4. Community composition of marine microbial communities based on metagenomes.** The top 10 bacterial and archaeal orders with highest mean relative abundance across all samples are shown. Relative abundances are based on the assembly and classification of the 16S rRNA gene by PhyloFlash. PPB = Port Phillip Bay, SML = Surface microlayer, STW = Subtropical waters, SAW = Subantarctic waters.

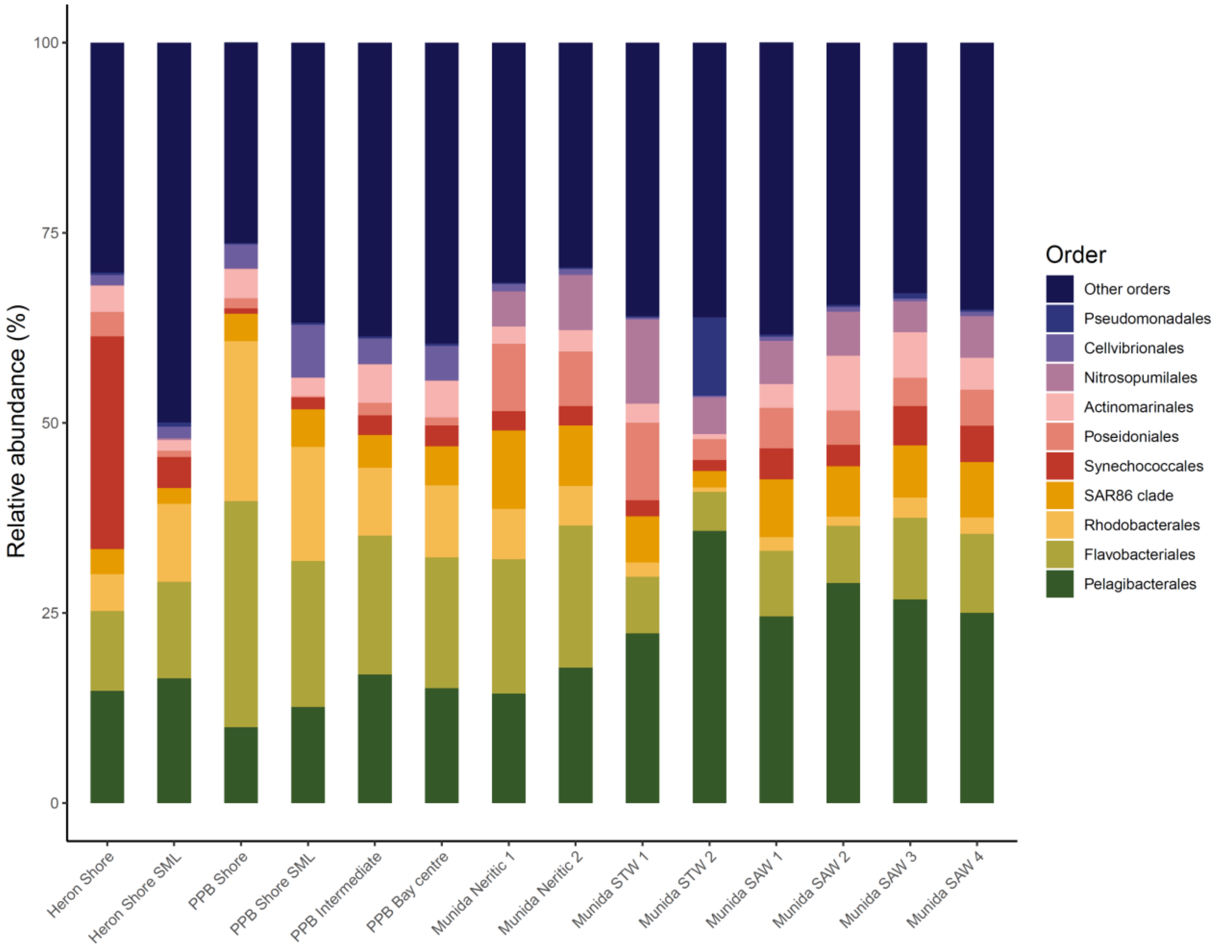

**Figure S5. Linear correlation between trace gas oxidiser abundance and trace gas oxidation rates.** $H_2$ and CO oxidizer abundance are based on the proportion of bacteria encoding $H_2$-uptake hydrogenases and form I CO dehydrogenases in the metagenomes. $H_2$ and CO oxidation rates are calculated based on bulk oxidation rates at measured *in situ* concentrations as per **Table S1**.

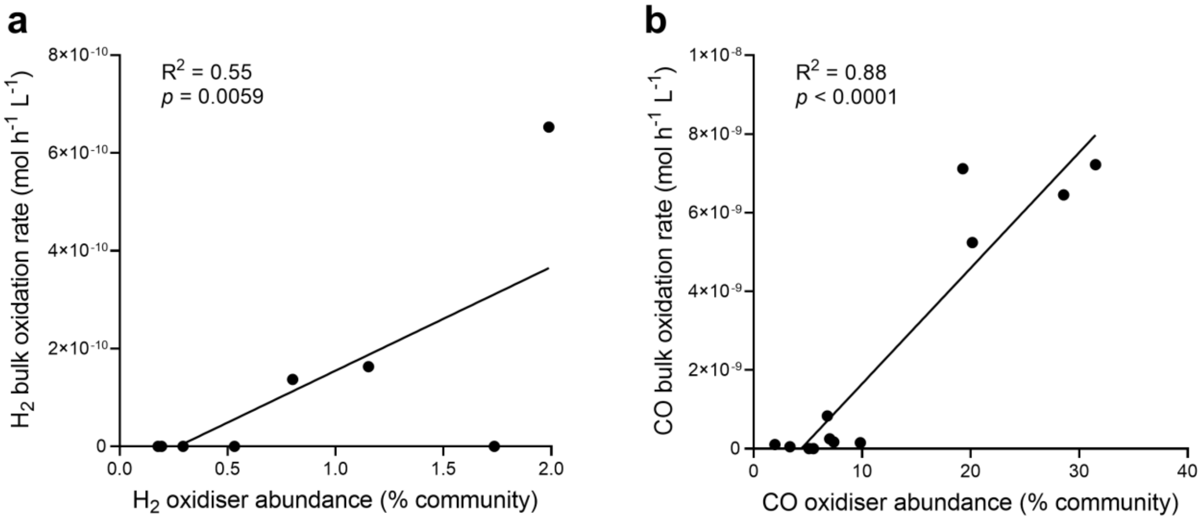
